# Pleural macrophages translocate to the lung during infection to promote improved influenza outcomes

**DOI:** 10.1101/2022.05.25.493482

**Authors:** James P Stumpff, Sang Yong Kim, Adriana Forero, Andrew Nishida, Yael Steuerman, Irit Gat-Viks, Meera G Nair, Juliet Morrison

## Abstract

Seasonal influenza results in 3 to 5 million cases of severe disease and 250,000 to 500,000 deaths annually. Macrophages have been implicated in both the resolution and progression of the disease, but the drivers of these outcomes are poorly understood. We probed mouse lung transcriptomic datasets using the Digital Cell Quantifier algorithm to predict immune cell subsets that correlated with mild or severe influenza A virus (IAV) infection outcomes. We identified a novel lung macrophage population that transcriptionally resembled small serosal cavity macrophages and correlated with mild disease. Until now, the study of serosal macrophage translocation in the context of infections has been neglected. Here, we show that pleural macrophages (PMs) migrate from the pleural cavity to the lung after infection with pH1N1 A/California/04/2009 IAV. We found that the depletion of PMs increased morbidity and pulmonary inflammation. There were increased proinflammatory cytokines in the pleural cavity and an influx of neutrophils within the lung. Our results show PMs are recruited to the lung during IAV infection and contribute to recovery from influenza. This study expands our knowledge of PM plasticity and provides a new source of lung macrophages independent of monocyte recruitment and local proliferation.

**GRAPHICAL ABSTRACT:** 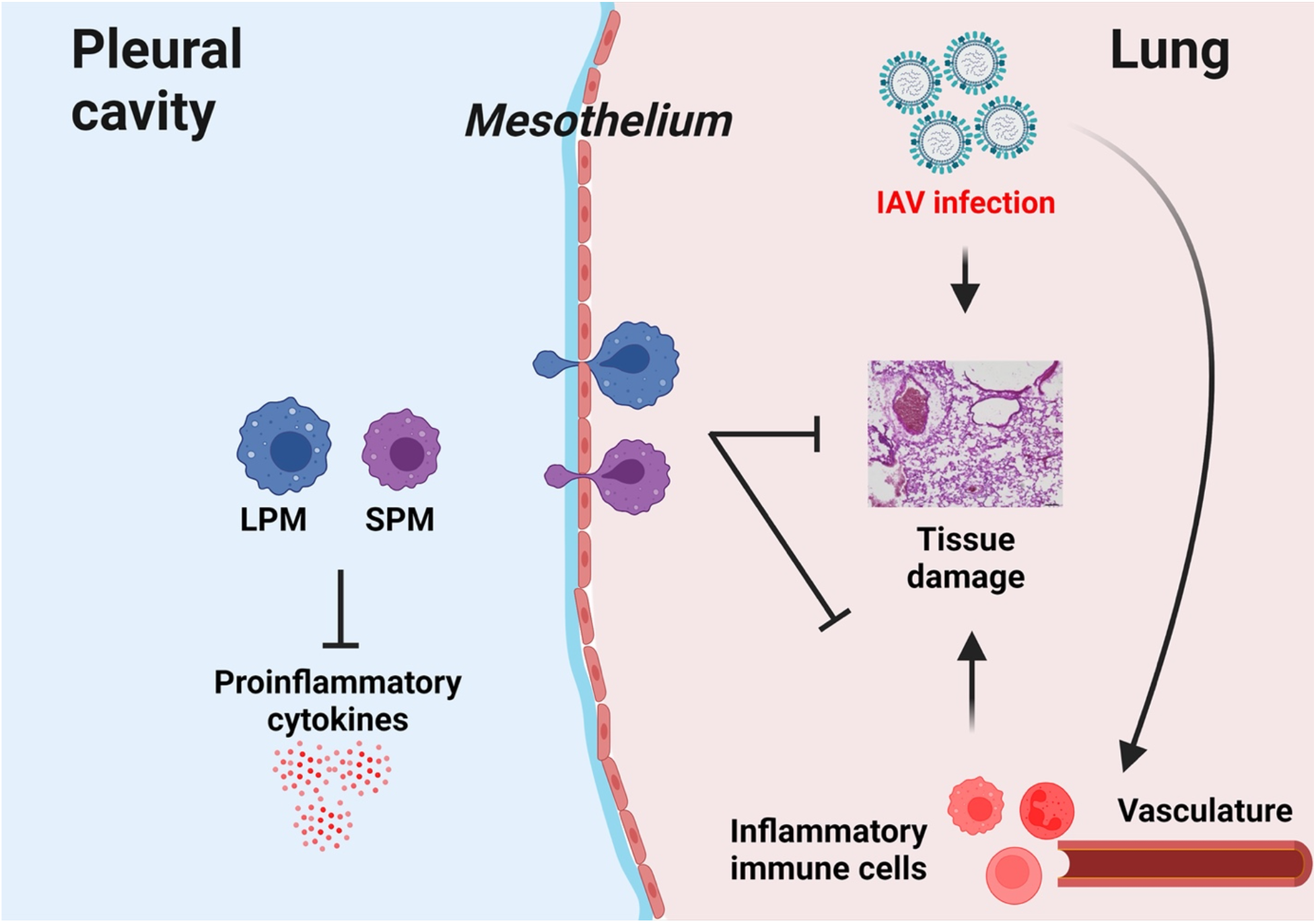

## INTRODUCTION

IAV is responsible for seasonal epidemics and several pandemics that arose from a lack of immunity and human-to-human transmission (1). Despite current vaccine strategies, IAV remains a major public health concern. Patient outcomes of IAV infections depend on the delicate balance between immune protection and immunopathology that is orchestrated by innate immune responses and subsequent adaptive immunity (2). Further investigation into IAV outcomes is needed to understand the resolution of viral clearance and restoration of pulmonary homeostasis.

The host response to infection is an important determinant of influenza outcomes (3-7). For example, severe influenza outcomes are associated with high levels of proinflammatory cytokines and leukocytes in the lung (5, 7, 8). Patients hospitalized with severe seasonal influenza infections have a sustained increase in monocytes (9), and patients with severe avian influenza have elevated levels of inflammatory cytokines in their acute-phase sera (10-12). Infection with highly pathogenic IAVs such as the 1918 virus and avian H5N1 virus leads to a massive recruitment of neutrophils and inflammatory macrophages to the lungs of mice (3, 4). Depending on the virus strain, mice may develop progressive pneumonia characterized by extensive neutrophilia, hypercytokinemia, pulmonary edema, and reductions in alveolar gas exchange that are reminiscent of acute respiratory distress (ARDS) in human patients (13-16). We previously identified lung transcriptomic signatures that distinguished mild and severe influenza outcomes in BALB/c mice infected with different IAV strains (17). A three-pronged lung signature consisting of decreased expression of lipid metabolism and coagulation genes and increased expression of proinflammatory cytokine genes had developed in mice that had succumbed to infection, while a signature of increased expression of lipid metabolism and coagulation genes and lower expression of proinflammatory cytokine genes had developed in mice that recovered from infection (17).

Serous membranes support and protect the internal organs of all vertebrate animals. Each serous membrane consists of two layers separated by a thin, fluid-filled serosal cavity. The serosal cavity that envelops the lungs is called the pleural cavity, while the cavities that surround the abdominal organs and the heart are known as the peritoneal cavity and the pericardial cavity, respectively. Serosal cavities contain multiple immune cells including innate B cells and T cells, but macrophages are the dominant cell population. Serosal macrophages are divided into small and large macrophages based on their cell size and surface marker expression. Small serosal macrophages are MHCII^+^F4/80^-^ and constitute ∼10% of serosal macrophages, while large serosal macrophages are MHCII^-^F4/80^+^ and comprise ∼90% of serosal macrophages (18-20). Serosal macrophages have been implicated in organ health. For example, postoperative gastrointestinal dysmotility can be ameliorated in mice by inhibiting peritoneal macrophage functions (21). Furthermore, large peritoneal macrophages enter the liver to promote wound healing in mouse models of sterile liver damage and dextran sulfate sodium (DSS)-induced intestinal colitis (22, 23), while large pericardial macrophages enter the heart to improve immune responses after myocardial infarction (24). However, small serosal macrophages have not been studied in diseases of the visceral organs, and serosal macrophages have never been studied in the context of viral infection.

In this manuscript, we use a systems biology approach as well as traditional “wet lab” techniques to identify a new lung macrophage population that originates in the pleural cavity and promotes recovery from influenza. To achieve this, we combined lung transcriptomic datasets to identify and confirm transcriptomic signatures that distinguish mild and severe influenza outcomes in mice. We then used a tissue deconvolution algorithm known as Digital Cell Quantifier (DCQ) to convert lung transcriptomic data into predictions of immune cell changes that precede different disease outcomes (25). We found that DCQ accurately predicted known cell population dynamics that occurred during influenza infection *in vivo*, and further predicted a lung cell population that transcriptionally resembled small serosal macrophages and whose numbers positively correlated with recovery from influenza.

We then used flow cytometry and microscopy to show that fluorescently-labeled PMs migrate from the pleural cavity into the lung after infection with a seasonal influenza virus strain, A/California/04/2009 (Cal09), after viral clearance has occurred and recovery has been initiated. We further show that depleting PMs leads to increased virus-induced weight loss and a longer recovery time. In addition, PM depletion causes increased inflammatory cytokine levels in the pleural cavity and increased neutrophil infiltration in the lung.

To our knowledge, we are the first to show that PMs translocate to the lung during IAV infection, and that PMs are important for the resolution of IAV-induced lung disease. We demonstrate the utility of our systems approach for discovering of immune cells subsets that correlate with mild and severe disease outcomes. Furthermore, our findings position the pleural cavity as an important contributor to lung homeostasis and the host response to pneumonia.

## RESULTS

### Host response differences in expression of inflammatory, metabolic, cell cycle and tissue repair genes distinguish influenza disease outcomes

Previously, we identified a gene expression signature in the mouse lung that could distinguish severe and mild influenza (17). We sought to expand our gene expression analysis to a wider range of influenza disease outcomes by including intermediate disease outcomes such as moderate weight loss and severe weight loss with subsequent recovery. Therefore, we integrated transcriptional data from our study (17) with data from an independent study of similar design (26). A description of the two studies and their combined weight loss outcomes are shown (**Figures 1A and 1B**). We restricted our analysis to those differentially-expressed (DE) genes that had a log fold change of 2 or more with an adjusted p-value cut-off of 0.05. When DE genes were clustered based on their biweight midcorrelation (bicor) across samples, six gene expression modules were identified and assigned unique colors (**Figure 1C**). We then used Ingenuity Pathway analysis (IPA) to assign functional categories to the genes within each module. The red module was enriched for genes in inflammation-associated pathways such as “Granulocyte adhesion and diapedesis” and “Crosstalk between dendritic cells and natural killer cells”. The increased induction of red module transcripts was associated with increased weight loss. The orange module, which had a unique expression pattern, was enriched for genes involved in lipid metabolism (“LXR/RXR activation” and “FXR/RXR activation” pathways) and coagulation. Upregulation of genes in this module on any day post infection in dataset GSE36328 or on day 3 post infection in dataset GSE54048 was associated with survival. Thus, expression patterns of genes within the red and orange modules support our previous observation of perturbations in inflammation, lipid metabolism, and coagulation signaling gene expression (17).

**Figure 1.**
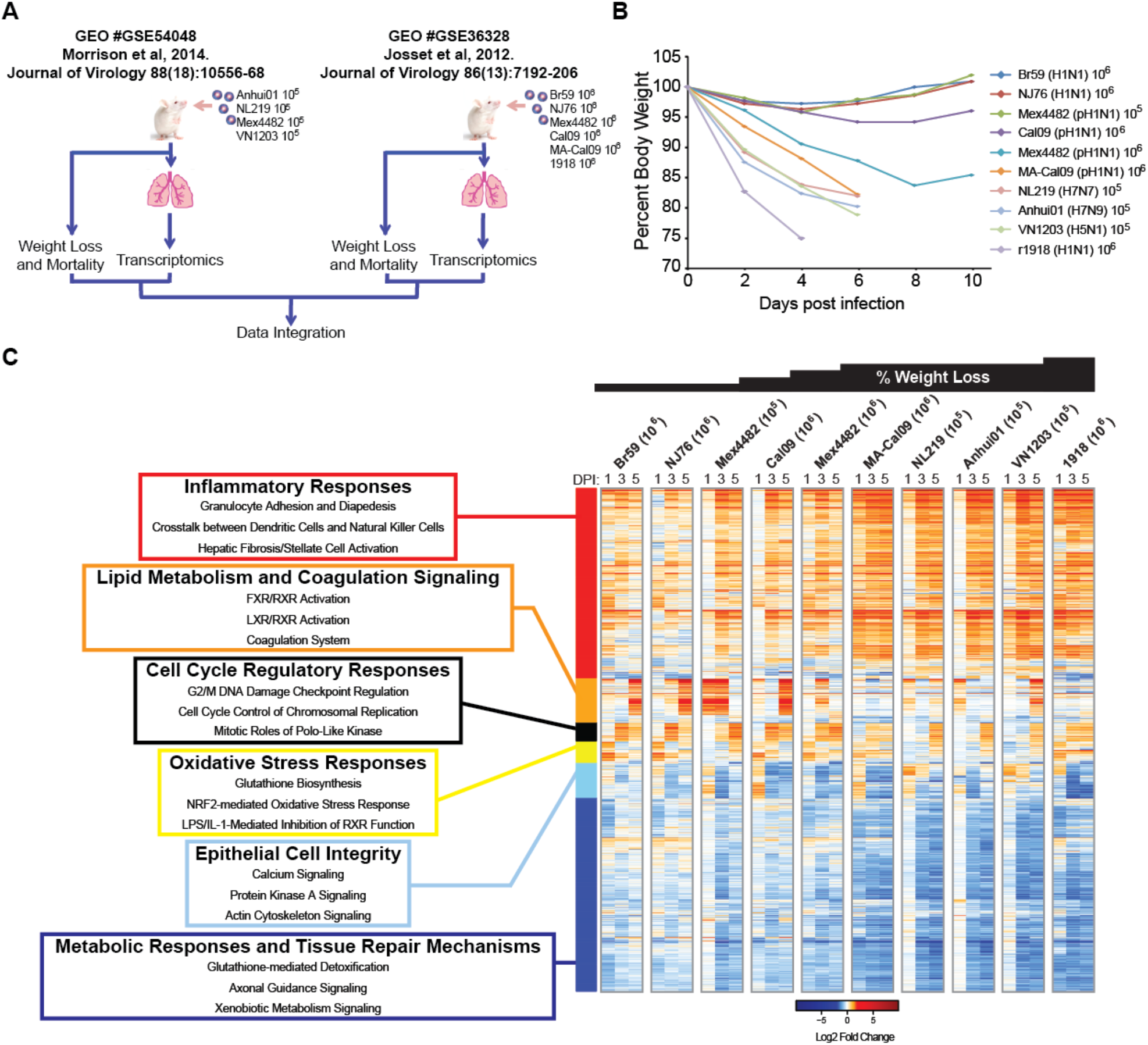
Different influenza disease outcomes are distinguished by host response differences in expression of inflammatory, metabolic, cell cycle and tissue repair genes. (A) Schematic showing the integration of BALB/C mouse data from GSE54048 and GSE36328. Experimental data were combined to produce (B) a combined weight loss dataset and a combined transcriptional dataset. (C) Hierarchical clustering of differential gene expression in murine lungs infected with influenza virus. Biweight midcorrelation clustering of 6012 genes that were found to be differentially expressed in any one condition (infection and time). Heatmap represents average gene expression intensity. Genes shown in red were upregulated and genes shown in blue were downregulated relative to uninfected lungs.

We then characterized the genes in the other four expression modules. The yellow module, which contained genes involved in oxidative stress responses, did not have an obvious pattern that related to weight loss or mortality. The sky-blue module, which contained calcium and actin cytoskeleton signaling genes, also lacked a pattern with regards to weight loss and mortality. Interestingly, upregulation of black module genes on day 3 post infection was associated with infection by H1N1 viruses, but this upregulation was unrelated to weight loss or mortality. The black module contained mitosis and cell cycle control genes. The downregulation of genes in the dark-blue module, which was enriched for tissue repair mechanisms and metabolic response genes, was associated with increased weight loss. We hypothesized that this signature resulted from differential activation or infiltration of immune cells in the lungs of mice that recovered versus those that succumbed to IAV infection.

### Several immune cell types are predicted to correlate with influenza disease severity

To identify immune cell populations that were potentially associated with the weight loss and mortality outcomes, we employed a tissue deconvolution method known as DCQ. The DCQ algorithm compares the gene expression profiles from 207 different immune cells with whole organ transcriptional data to predict the quantities of immune cells within a complex organ (25). This method utilizes a panel of genes encoding cell surface markers that are commonly used for flow cytometry, and whose transcript and protein levels are concordant (25). We took the populations measured by DCQ and used linear regression to identify the cell populations whose numbers were most highly associated with weight loss in infected animals. We identified 26 cell types that were positively or negatively correlated with influenza outcomes (**Figure 2**). Loss of stem cell populations as well as lymphoid cells such as immature B and T cells and effector CD8^+^ T cells was associated with increased weight loss. An increase in monocytes, plasmacytoid dendritic cells (pDCs) and granulocytes was associated with increased weight loss. Though increases in several conventional dendritic cell (cDC) and MF populations were associated with increased weight loss and death, the presence of other cDC and MF populations were associated with mild disease and recovery. For example, decreases in CD103^+^ cDCs (DC.103^+^11B^-^.LU) were associated with increased morbidity and mortality. The presence of cells resembling MHCII^+^F4/80^lo^ peritoneal MFs (MF.II^+^480^lo^.PC) were associated with mild disease.

**Figure 2.**
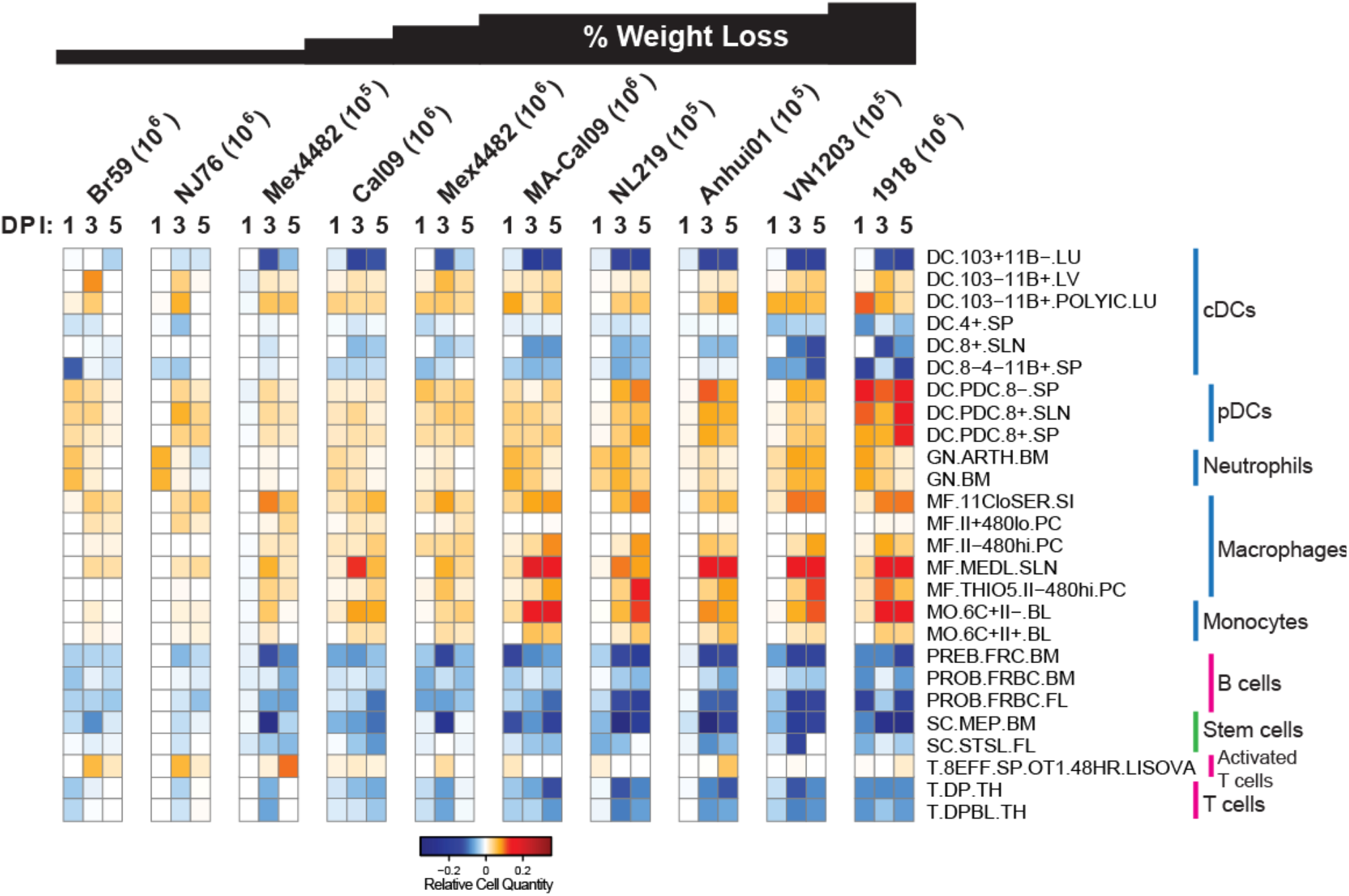
Digital cell quantifier identified immune cell subsets that predict disease severity or recovery across independent experiments. We surveyed the *in vivo* dynamics across time and viral strains using digital cell quantifier (DCQ; http://dcq.tau.ac.il). Linear regression models revealed distinct immune cell populations predicted to drive disease morbidity as defined by weight loss following influenza virus infection. The heatmap represents the relative quantity of cell types with significant relationships (p < 0.05) between that cell type on at least one day to the weights on at least one day after filtering for observations where data for at least 8 samples were available.

### Immune cell predictions are conserved across multiple transcriptomic datasets

To further bolster the tissue deconvolution predictions, we subjected two additional microarray datasets to DCQ. In Shoemaker *et al*. (GSE63786), C57BL/6 mice were infected with 10^5^ PFU of Cal09 or VN1203, and monitored over the course of 7 days (27) (**Figure S1A**). Lungs from VN1203-infected mice were found to have higher viral loads and more pathology than lungs from Cal09-infected mice (27). In McDermott *et al*. (GSE33263), C57BL/6 mice were infected with 10^2^, 10^3^ or 10^4^ PFU VN1203 (28) (**Figure S1B**). Higher inoculation titers led to increased weight loss and mortality (28). When GSE63786 and GSE33263 were run through the DCQ algorithm, we found that the 26 cell types identified from the BALB/c datasets (**Figure 2**) largely showed a similar pattern in the C57BL/6 data (**Figures S1C and S1D**). As with the BALB/c mice, an increase in monocytes and granulocytes was associated with increased tissue pathology and weight loss in C57BL/6 mice (**Figures S1C and S1D**). Loss of stem cell populations, immature B and T cells and effector CD8^+^ T cells was also associated with increased weight loss. Again, the presence of cells resembling MHCII^+^F4/80^lo^ peritoneal macrophages was associated with mild disease (**Figures S1C and S1D**). The only prediction that did not hold across the 4 datasets was that for pDC populations. Higher pDC numbers were associated with severe disease in BALB/c mice (**Figure 2**) but were associated with mild disease in C57BL/6 mice (**Figure S1**).

### Flow cytometry validates DCQ predictions

DCQ accurately identified cell population dynamics that have been shown to occur during IAV infection *in vivo* (**Figures 2** and **S1**). For example, CD103^+^CD11b^-^ dendritic cell numbers decrease post infection, modeling what occurs *in vivo* when they exit the lung and traffic to the draining lymph nodes to present antigen to CD8^+^ T cells (29, 30).

To further emphasize the value of our approach, we conducted *in vivo* experiments to confirm some of the predictions. BALB/c mice were infected intranasally with 10^4^ pH1N1 A/California/09 (Cal09) virus to induce mild disease or 10^4^ H1N1 A/Puerto Rico/1934 (PR8) virus to induce severe disease. Control mice received PBS intranasally. We isolated, stained, and subjected lung cells to flow cytometry on day 3 post infection. We validated the prediction that more neutrophils and Ly6C^+^ monocytes were recruited to the lung during severe disease (**Figure S2**) as has been described before (4, 9, 14, 31, 32).

### Influenza virus infection promotes the recruitment of pleural macrophages to the lung

Our data thus far supported the idea of a MHCII^+^F4/80^lo^ macrophage population that originates in a serosal cavity and is present in the lungs of mice that recover from influenza. Since the macrophage populations of the pleural and peritoneal cavities are analogous (18, 20), and the pleural cavity envelopes the lung, we hypothesized that the MHCII^+^F4/80^lo^ lung macrophages predicted by DCQ originated in the pleural cavity. Though CD11b, CD115, F4/80 and MHCII are sufficient for distinguishing the two pleural macrophage (PM) populations, these markers are insufficient for distinguishing the various macrophage populations in the lung. To circumvent issues with lung macrophage identification, we focused instead on the potential origin of the novel lung population. PMs were labeled *in vivo* by injecting a red phagocyte-specific dye (PKH26PCL) into the pleural cavities of mice 1 day prior to infection. When mice were infected intranasally with 10^2^ PFU of Cal09, PKH26PCL-labelled cells accumulated in the lung (**Figure 3**).

**Figure 3.**
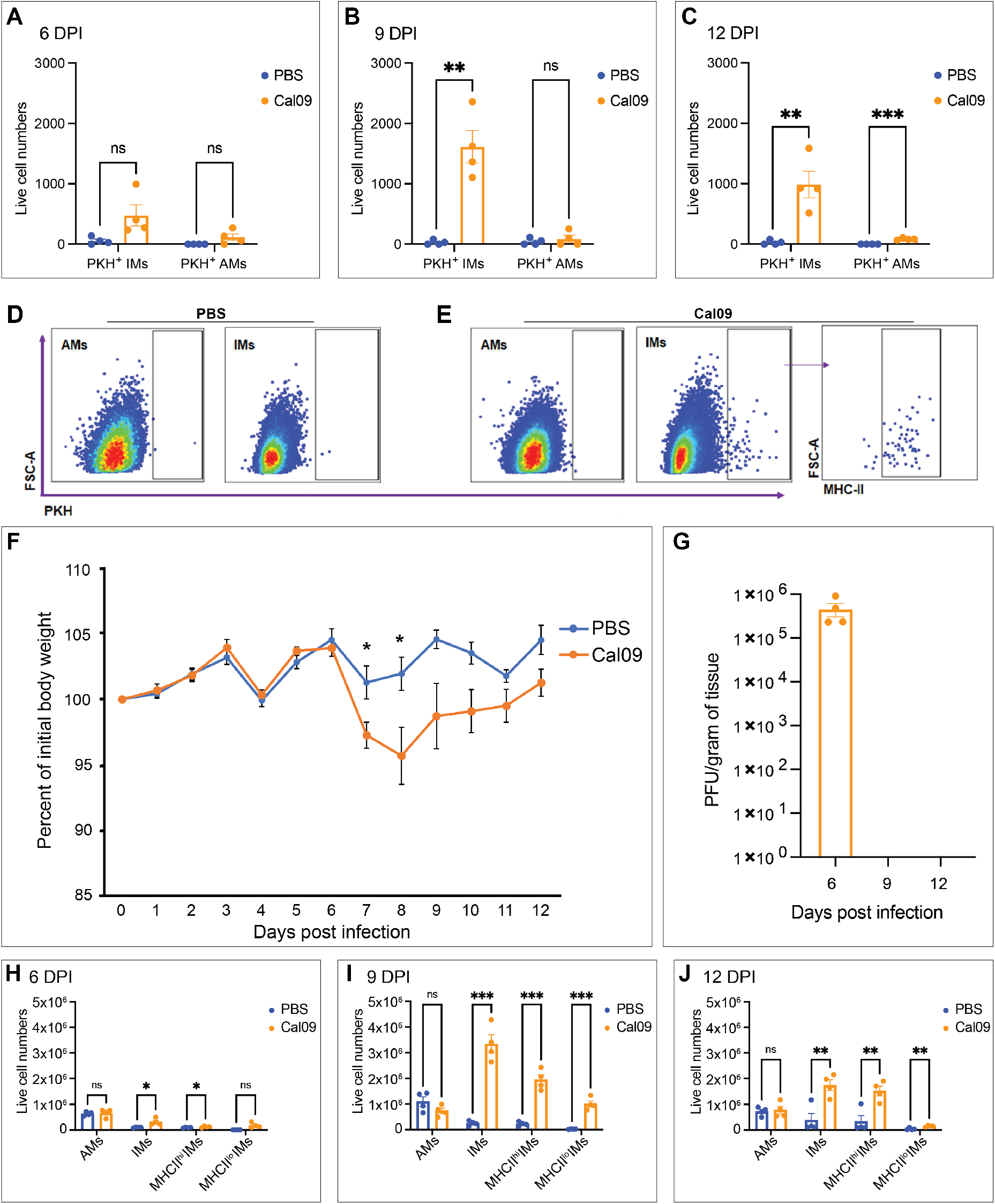
Influenza virus infection promotes the recruitment of pleural macrophages to the lung. BALB/c mice were intrapleurally injected with PKH26PCL dye one day before they were intranasally infected with 10^2^ Cal09 virus or mock-infected with PBS as a control. (A-C) Quantification of live cell numbers of PKH^+^ AMs and IMs in Cal09- or mock-infected mice. (D) Representative flow plots of PBS control mice. (E) Representative flow plots of Cal09 infected mice. (F) Representative growth curve of Cal09-infected versus PBS control mice. (G) Virus titers from lungs of mice in (F). (H-J) Quantification of live cell numbers of AMs and IMs in Cal09- or mock-infected mice. Data shown as mean ± SEM (n=4 from each group, *P<0.05, **P<0.01, ***P<0.001, Student’s t-test). AM = alveolar macrophage (MerTK^+^CD64^+^SiglecF^+^CD11b^-^); IM = interstitial macrophage (MerTK^+^CD64^+^CD11b^+^SiglecF^-^).

Translocated PMs were detected in the lungs on days 6, 9, and 12 post infection, but more accumulation of the PKH^+^ PMs occurred on 9 and 12 days post infection when compared to the uninfected controls (**Figures 3A-E**). Flow cytometry gating strategies to distinguish pleural and lung macrophage subpopulations are outlined in **Figures S3** and **S4**.

PKH^+^ cells in the lung were CD64^+^MerTK^+^SiglecF^-^CD11b^+^, which are phenotypically like interstitial macrophages (IMs) (**Figure 3E**). A smaller pool of PMs did phenotypically resemble CD64^+^MerTK^+^SiglecF^+^CD11b^-^ alveolar macrophages (AMs) (**Figure 3E**). Heterogeneity amongst IMs has been researched in multiple studies (33-35). One way of distinguishing them is based on their expression on MHCII. We found that the majority of PKH^+^ lung cells resembled MHCII^+^ IMs (**Figure 3E)**. Accumulation of this MHCII^+^CD64^+^MerTK^+^SiglecF^-^CD11b^+^ population that had originated in the pleural cavity only occurred once animals began to regain weight and after they had cleared virus from their lungs (**Figures 3F-G**). We also observed an overall increase in the total number of IMs in the lung during IAV infection (**Figures 3H-J**).

To establish the location of the PKH^+^ PMs in the lung, frozen lung sections were immunostained and imaged by fluorescence microscopy 9 days post infection (**Figure 4A**). PKH^+^ PMs in the lung are detected near the mesothelium (shown by the white dotted line) and within regions of dense DAPI signal (**Figure 4A)**. No PKH^+^ PMs were detected in the PBS mock-infected control group. To determine whether the migration of PMs occurs through the vasculature, immune cells were isolated from blood on 6, 9, and 12 days post infection. Flow cytometric analysis showed no PKH^+^ PMs in the blood from either the infected or the control group indicating that the labeled PMs had trafficked through the mesothelium (**Figure 4B)**.

**Figure 4.**
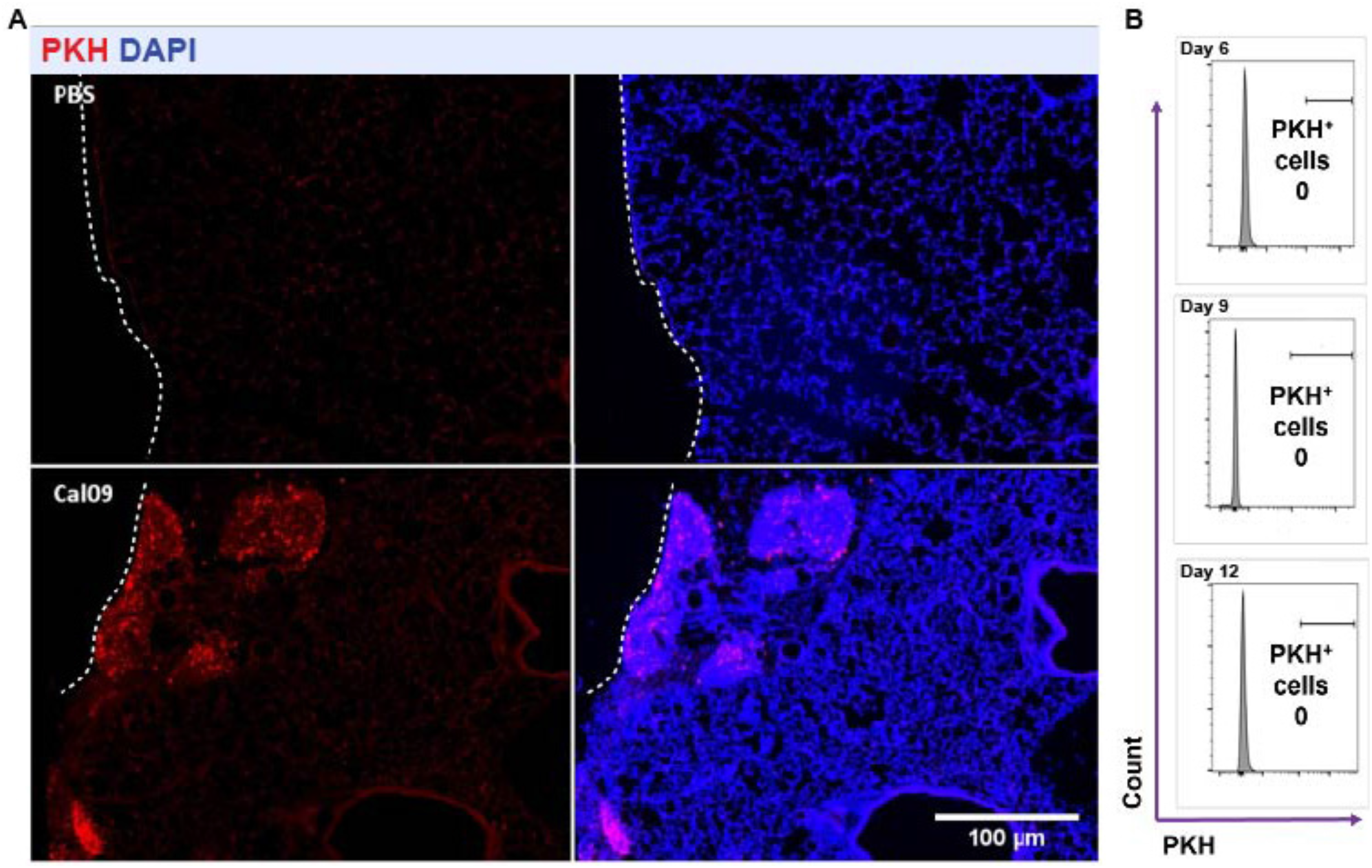
PMs localize near the mesothelium. (A) Representative fluorescent images of BALB/c naïve and Cal09-infected lungs harvested 9 days post infection. PMs were labeled *in vivo* with PKH26PCL dye (red) and counterstained with DAPI (blue). The mesothelium is depicted by the white dotted line (n = 3-7 from each group). (B) Flow cytometry of blood harvested from mice that had been intrapleurally injected with PKH26PCL dye then infected with Cal09 on days 6, 9 and 12 post infection.

### Pleural macrophage depletion leads to increased weight loss and slower recovery from IAV infection

Previous studies have shown evidence for a role of cavity macrophages in models of liver, heart and intestinal injury, showcasing differences in recruitment, wound repair, and weight loss (22-24, 36). Another study suggested a role for PMs in bacterial clearance in bacterial pneumonia (37). However, no study to date has investigated the role of PMs in viral infection. To test whether PMs affect influenza outcomes, we depleted PMs by injecting clodronate liposomes (CLL) into the pleural cavities of mice 1 day prior to infection with Cal09 virus. Flow cytometry confirmed PMs were depleted 24 hours after injection (**Figure S5**), and up to 14 days after injection. PM-depleted mice lost more weight compared to the PBS liposomes (PBSL)-injected control mice (**Figure 5A**). However, we noted no statistical differences in H&E-stained lungs (**Figure S6A**) that had been scored for airway thickening, alveolar destruction, and vascular inflammation on 9 days post infection (**Figure S6B**). Lungs viral loads were unaffected by PM depletion (**Figure 5B**).

**Figure 5.**
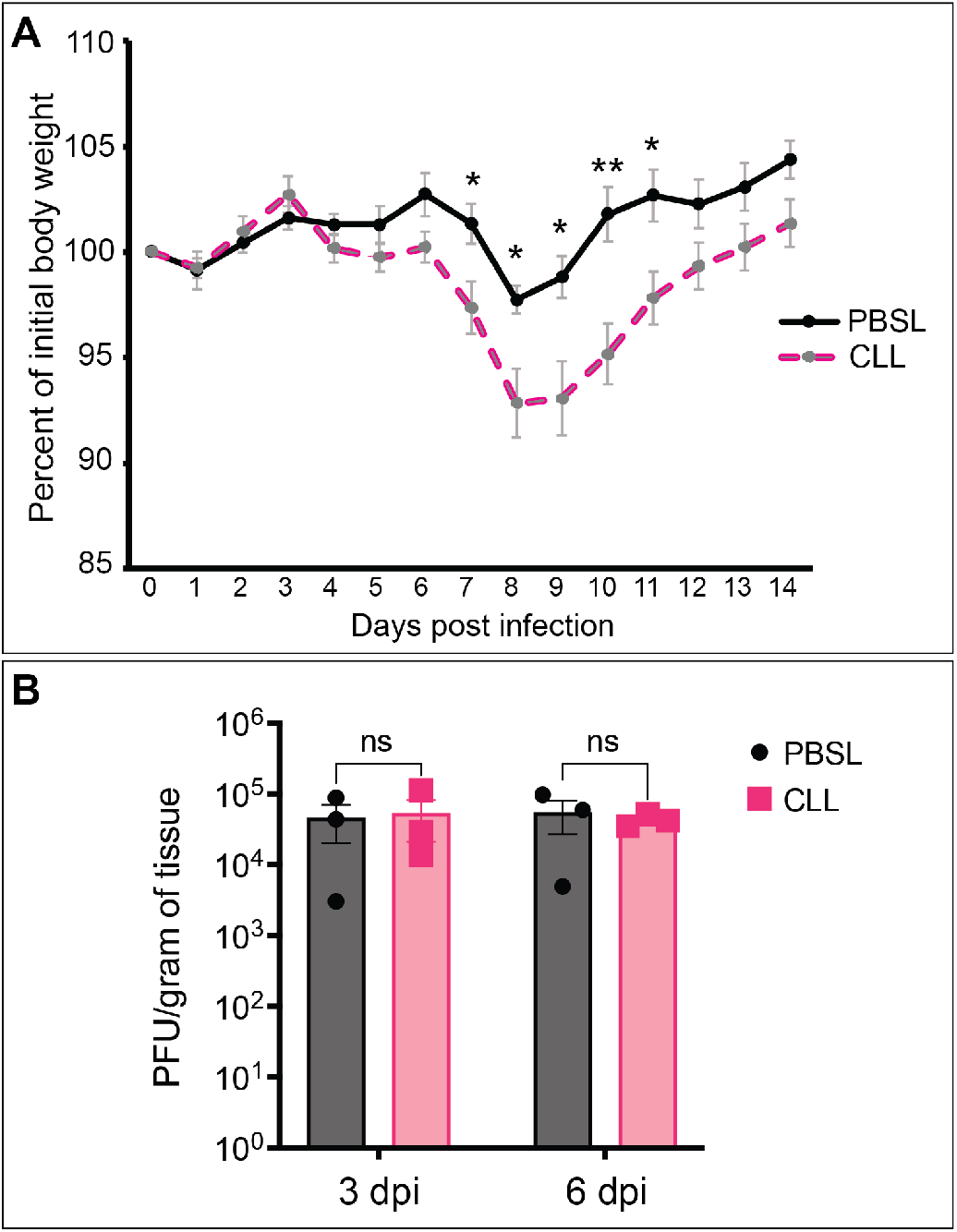
PM depletion increases disease severity without impacting viral titers. BALB/c mice received an intrapleural injection of CLL or PBSL one day before infection with 10^2^ PFU of Cal09. (A) Weight loss was tracked over 14 days post infection in both CLL-injected and PBSL-injected groups (n=10 per group). (B) Virus titers from lungs of mice (n=3 per group). Data shown as mean ± SEM (*P<0.05, **P<0.01, ***P<0.001, ****P<0.0001, Student’s t-test).

Since PM depletion led to more IAV-induced weight loss, we asked whether we would see increased inflammation in the lungs of PM-depleted mice. By measuring the number of CD45^+^ cells in the lung, we found that the number of total leukocytes in the lungs of PM-deficient and PM-sufficient mice were the same (**Figure 6A**). However, when we looked at individual leukocyte populations, we found that CLL-treated mice had a significantly higher percentage of neutrophils than PBSL-treated mice did on day 9 post infection (**Figure 6B**). Our flow cytometry gating strategy for distinguishing lung leukocyte populations is outlined in **Figure S7**. There were fewer AMs in CLL-treated mice (**Figure 6C**). There were also fewer interstitial macrophages in CLL-treated mice albeit not statistically significant (**Figure 6B**). No significant differences were observed in monocyte and dendritic cell subsets (**Figure 6D-E**). Additionally, PM depletion led to increased pleural cavity inflammation indicated by increased proinflammatory cytokines: interferon-gamma (IFN-γ), tumor necrosis factor alpha (TNF-α) and monocyte chemoattractant protein-1 (MCP-1) (**Figure 7A**). However, no proinflammatory cytokines were increased in the lung between the infected groups (**Figure 7B**) though PM-depletion caused an increase in serum IFN-γ levels in response to lung IAV infection at that time point (**Figure 7C**).

**Figure 6.**
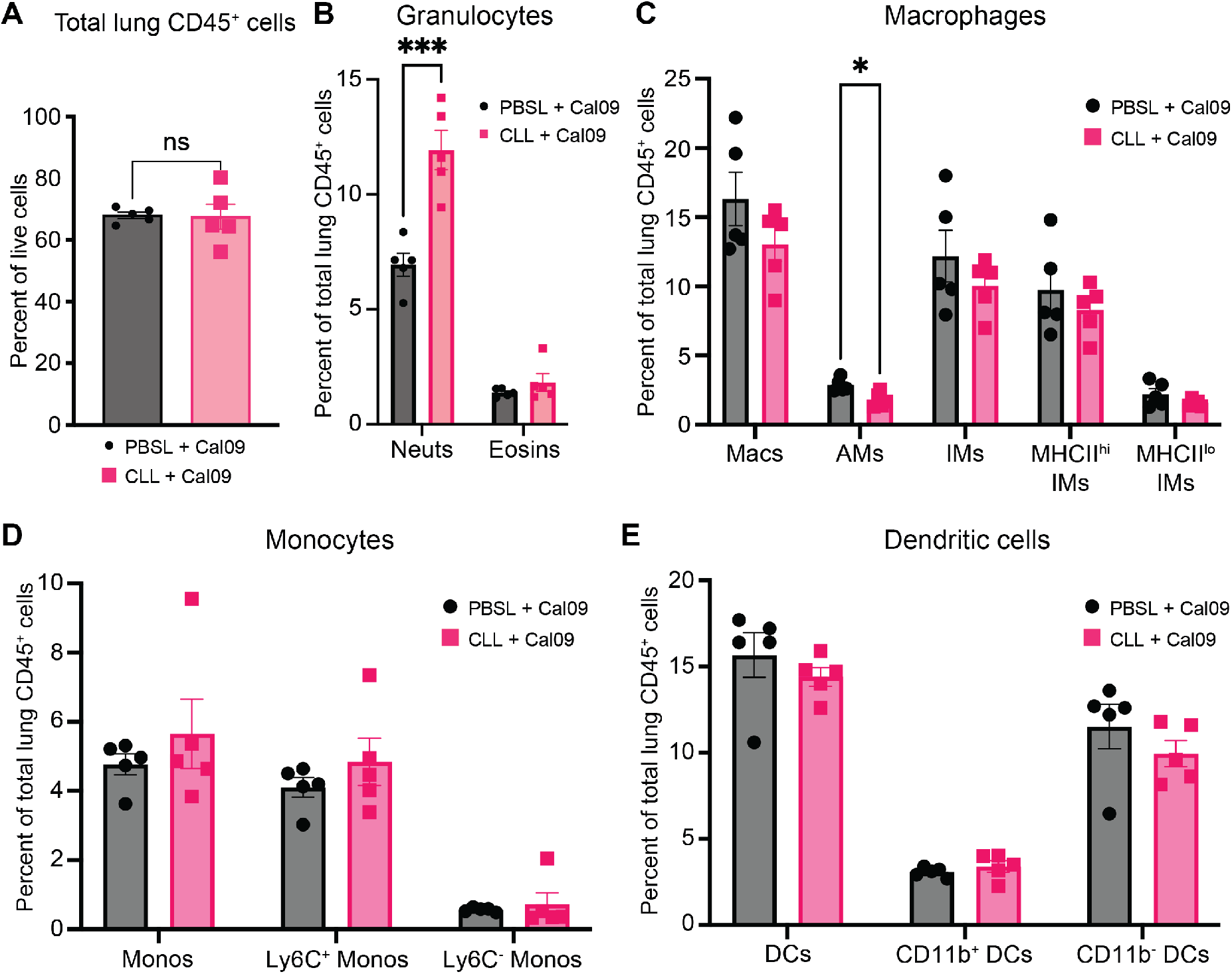
PM depletion increases neutrophil infiltration in the lung on day 9 post infection. BALB/c mice received an intrapleural injection of CLL or PBSL one day before infection with 10^2^ PFU of Cal09, and lungs were isolated for flow cytometry on day 9 post infection. (A) Flow cytometric analysis of lung leukocytes (CD45^+^), (B) neutrophils (Ly6G^+^CD11b^+^) and eosinophils (CD11b^+^SiglecF^+^), (C) *left to right*, macrophages (CD64^+^MerTK^+^) were divided into alveolar macrophages (SiglecF^+^CD11b^-^) and interstitial macrophages (CD11b^+^SiglecF^-^), which were further divided into two IM subsets (MHCII^hi/lo^), (D) monocytes (CD64^lo^) and two subpopulations (Ly6C^+/-^), (E) dendritic cells (CD11c^+^MHCII^+^CD24^+^) and two subpopulations (CD11b^+/-^). Data shown as mean ± SEM (n=5 from each group, *P<0.05, **P<0.01, ***P<0.001, ****P<0.0001, Student’s t-test). Neuts = neutrophils, eosins = eosinophils, macs = macrophages, AMs = alveolar macrophages, IMs = interstitial macrophages, monos = monocytes, DCs = dendritic cells

**Figure 7.**
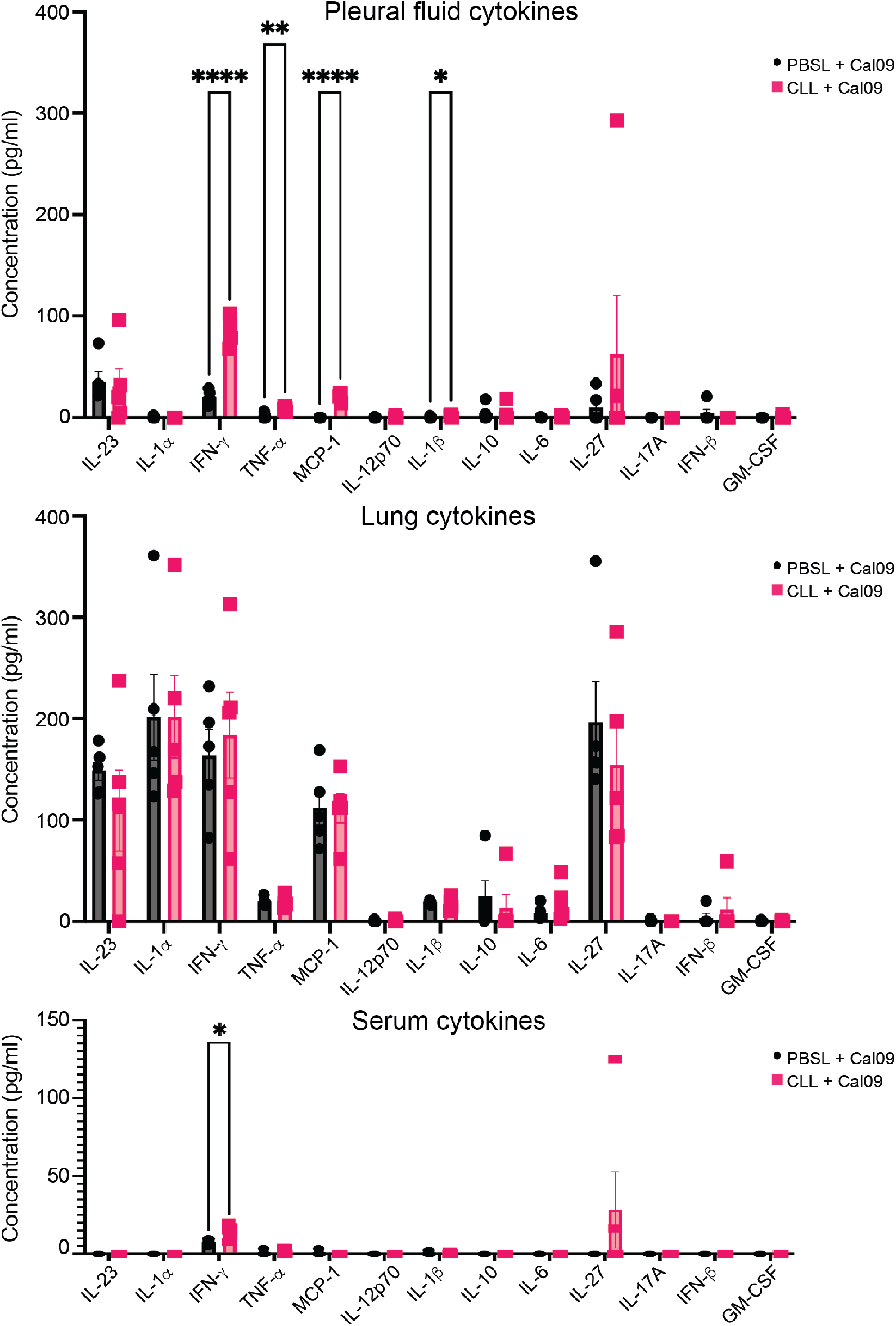
PM depletion leads to more proinflammatory cytokines in the pleural cavity on day 9 post infection. BALB/c mice received an intrapleural injection of CLL or PBSL one day before infection with 10^2^ PFU of Cal09, and pleural fluid, lungs and sera were isolated on day 9 post infection. Cytokines from (A) pleural fluid, (B) lung homogenate, and (C) serum. Data shown as mean ± SEM (n=5 from each group, *P<0.05, **P<0.01, ***P<0.001, ****P<0.0001, Student’s t-test)

## DISCUSSION

The impact of serosal macrophages on visceral organs has been an understudied area of research. However, a few key studies have described migration of serosal macrophages into visceral organs. These foundational studies focused on sterile injury or inflammation of the liver, heart, and intestine (22-24). Here, we identified a previously unrecognized role for PMs in influenza. We found that IAV infection triggers the recruitment of mature macrophages from the pleural cavity, across the mesothelial layer, and into the lung. PMs are recruited after the clearance of viral infection and when restoration of homeostasis is critical. We observed this by labeling PMs via intrapleural injection of PKH26PCL prior to IAV infection then measuring their translocation using flow cytometry and immunofluorescence (**Figures 3** and **4**). A recent study defined one population of IMs as nerve- and airway-associated macrophages, which express MHCII and proliferate rapidly after IAV infection (38). Our data are supportive of this; we observed a robust increase in the numbers of IMs, most of which are an MHCII^hi^ subpopulation that peaked at 9-days post infection. We show that PMs that translocate to the lung contribute to this MHCII^hi^ IM pool (**Figure 3**).

Our results differ from those of a recent study that described only surface accumulation of pleural and peritoneal macrophages after organ injury, and did not identify a role for serosal macrophages in tissue repair or regeneration (36). This difference may be explained by the timepoints used for observation. We visualized PM translocation at late timepoints−during the resolution phase−of a seasonal IAV infection model and showed that PMs affect disease severity. Ablating PMs one day prior to infection led to increased weight loss and neutrophil influx in the lungs of IAV-infected mice.

While we were preparing this manuscript, Bénard *et al*. described increased bacterial burden and mortality upon PM depletion in a mouse model of bacterial pneumonia (37). Unlike in Bénard *et al*., where bacterial burdens increased upon PM depletion and resulted in increased mortality, we saw increased disease severity without differences in viral titers between PM-depleted and control groups after IAV infection. Additionally, we showed that PMs traffic to the lung after IAV infection, while they did not report PM translocation.

An important feature of our study was the use of a systems approach to generate hypotheses. We identified signatures that distinguished mild and severe disease outcomes in mice by combining lung transcriptomic data from two IAV mouse infection studies. We then used tissue deconvolution and linear regression to screen immune populations from the ImmGen database that could be driving differences in infection outcomes. Though there is a great deal of heterogeneity amongst lung macrophage populations (19, 20, 33-35, 39), the DCQ algorithm accurately predicted the presence of a population resembling small serosal macrophages that accumulated only in the lungs of animals that survived IAV infection. This prediction was foundational and served as an important tool for identifying a novel immune cell population in our system. Nevertheless, there are limits to tissue deconvolution approaches. Many macrophage populations from different organs share similar phenotypic markers which make it difficult to predict the origin of immune cell populations. In our case, the uniqueness of the small peritoneal macrophage transcriptome in ImmGen was an asset to our predictions and subsequent analyses. As such, characterizing additional immune populations of the serosal cavities and other unique niches will be useful for future predictions.

PMs are important immunomodulators that impact the recovery of IAV-infected mice by decreasing pleural space inflammation and lung neutrophil infiltration (**Figure 6**). Tissue resident macrophages have been shown to “cloak” proinflammatory debris to contain neutrophil-driven tissue damage and inflammation (40). Furthermore, attenuation of neutrophil influx in IAV infection can improve survival without impacting viral titers (31). Thus, it is feasible that recruited PMs may play a role in masking damage signals to prevent neutrophil infiltration. Pulmonary inflammation can lead to pleural inflammation, which is associated with increased mortality in pneumonia patients and in patients hospitalized with pleural effusion (41, 42). IFN-γ levels were higher in both pleural fluid and sera of IAV-infected, PM-depleted mice (**Figure 7**). IFN-γ deficiency has been shown to decrease susceptibility to lethal infection by increasing activation of group II innate lymphoid cells (ILC2s) (43). We also saw increased MCP-1 in the pleural cavity, and this has been shown to contribute to pleurisy and pleural effusion in carrageenan-induced pleurisy (44).

The pleural cavity may serve as a reservoir for other immune cells that can migrate to the lung. B1a cells, another pleural cavity immune cell population, have important roles in bacterial pneumonia and were shown to migrate to the lung after LPS challenge (45). Furthermore, during IAV infection, pleural cavity B1a cells have important pulmonary responses and are suggested to migrate to infected lungs (46). Whether other pleural cavity cell populations can migrate to the lung has not been determined, but when we looked at other innate immune cell populations, we did not detect PKH^+^ cells.

The role of the pleural cavity in other viral infections and secondary bacterial infections remains poorly understood. However, meta-analyses of SARS-CoV-2-infected patients showed that pleural effusion was associated with poorer COVID-19 prognoses (47). Similar observations have been seen in patients infected with IAV and/or bacteria (41, 42, 48-50). Additionally, pleural space inflammation can provide protection against bacterial lung infection (37). Following viral infection, patients are left more vulnerable to subsequent pneumonia, but PMs may limit inflammation after infection and decrease susceptibility to pneumonia. There is therefore a need for more research into pleural cavity function in the context of lung infection. Altogether, we show that PMs migrate to the lung during IAV infection and play a role in limiting disease severity by modulating inflammatory responses both in the pleural cavity and in the lung. Selectively targeting PMs could serve as a strategy for treating severe influenza and other lung and pleural diseases. Future studies to understand how PM translocation and plasticity are regulated are warranted.

## METHODS

### Gene expression analysis

The primary transcriptomic data sets GEO Accession number GSE54048, GSE36328, GSE63786 and GSE33263 (17, 26-28) were extracted and quantile normalized using the “normalizeBetweenArrays” method available in the “limma” package of the R suite (51), and adjusted for batch effects using ComBat software (52). Expression across each sample was normalized to the average expression of study- and time-matched mocks. Differential gene expression following viral infection was determined by deriving the ratio of expression between the average gene expression of influenza virus-infected replicates to the average of time-matched mock-infected samples applying a linear model fit using the “limma” package (51). Criteria for differential expression (DE) were an absolute log-fold change of 2 and an adjusted *P* value of <0.05, calculated by Benjamini-Hochberg correction. Functional analysis of DE genes was done using Ingenuity Pathway Analysis (IPA; Ingenuity Systems) using a right-tailed Fisher exact test with a threshold of significance set at *P* value of 0.05.

### Computational measurement of immune cell subsets

Immune cell dynamics during the course of infection were surveyed using the Digital Cell Quantifier (DCQ) algorithm as previously described (25). Genome-wide gene expression was normalized for each infection relative to the average of the time-matched mock, and a log 2 transformation applied. Each gene expression entry was divided by the standard deviation across test samples. Analysis of relative cell quantity was run with three repeats and a lambda minimum of 0.2. For each of the 207 cell types in DCQ and their three time points on day 1, day 3, and day 5, we used a simple linear model to estimate the relationship between the sample’s expression for that cell type on that day to the weights on days 2, 4, 6, 8, 10, and 12.

### Viruses

pH1N1 A/California/04/2009 and H1N1 A/Puerto Rico/8/1934 viruses were a kind gift from Dr. Adolfo García-Sastre (Icahn School of Medicine at Mount Sinai). Virus stocks were propagated in 8-day-old embryonated eggs (Charles River Laboratories) and titrated by plaque assays on MDCK cells.

### Mouse experiments

Female BALB/c mice, 8 to 12 weeks old, were purchased from Jackson Laboratory (Bar Harbor, ME) and used for all mouse experiments.

Mice were injected intrapleurally, under isoflurane anesthesia, with 100 μl of 10μM PKH26PCL fluorescent dye (Sigma Aldrich) one day before virus infection. In other experiments, mice were injected intrapleurally, under isoflurane anesthesia, with 50 μl of clodronate liposomes or PBS control liposomes (Encapsula Nano Sciences) one day before virus infection. Mice were challenged intranasally, under isoflurane anesthesia, with 50 μl PBS containing 10^2^ or 10^4^ PFU of pH1N1 A/California/04/2009 or H1N1 A/PR/8/1934 viruses, or mock challenged with PBS alone. Lungs were collected and homogenized in 1 ml PBS and stored at −80 °C until virus titration. Viral titers were measured by plaque assays on MDCK cells.

### Tissue homogenization

Lungs were harvested at indicated timepoints and washed in PBS before being placed in 2-ml tubes containing 1 ml 1X PBS, 3% FBS, and 1mM EDTA, and one ceramic bead (MP Biomedicals). Lung tissue was homogenized with the FastPrep-24 bead beater at 7.00 speed/s twice for 20 seconds each then placed on ice for 5 min and repeated. After homogenization, samples were centrifuged at 21,300xg for 15 min. Supernatant was aliquoted into new tubes and stored at -80°C.

### Plaque assay

MDCK cells were seeded in 6-well culture plates and incubated at 37°C in 5% CO_2_ for 24 hours. 10-fold serial dilutions were prepared in a solution containing 1x PBS, 0.21% BSA, 1% Pen/Strep, and 1% Ca/Mg. Cells were washed once with PBS before adding 200 ml of the serial dilutions. Plates were incubated at 37°C and were rocked side-to-side and forward-to-back every 15 minutes to distribute virus inoculum over the monolayer of cells for 1 hour. TPCK-treated trypsin (1 mg/ml) was added at a 1:1000 ratio to a supplemented 2X DMEM (2X DMEM, 2% Pen/Strep, 0.42% BSA, 20 mM HEPES, 0.24% NaHCO_3_, 0.02% DEAE-Dextran) before mixing with 1.5% Oxoid Agar at a 1:1 ratio. 2 ml of the agar overlay was added to each well and allowed to cool for 15 min at room temperature before transferring to the 37°C incubator. Plates were incubated for 96 hours. After incubation, plates were fixed with 1 ml of a 3.7% formaldehyde solution and incubated for 10 minutes to neutralize infectious virus. Overlay was flicked out with a spatula and stained for 20 min with crystal violet (0.095% crystal violet, 2.8% ethanol, 19% methanol). Plates were rinsed in tap water and plaques were counted to determine viral titers.

### Cytokine and chemokine detection

Cytokines and chemokines from pleural fluid, serum, and lung were quantified using the LEGENDplex™ (Biolegend, 740150) mouse inflammation panel (13-plex). All samples were analyzed on a NovoCyte Quanteon and data was analyzed using LEGENDplex™ software (Biolegend).

### Preparation of cell suspensions, flow cytometry and cell sorting

For isolation of pleural cavity cells, the pleural cavity was washed twice with 500 μl sterile cold PBS. The fluid was centrifuged at 500xg for 5 min at 4°C and resuspended in RBC lysis buffer (Sigma, R7757) for 5 min at room temperature. Fluorescence-activated cell sorting (FACS) buffer (1x PBS, 3% FBS and 2 mM EDTA) was added to stop the lysis and the fluid was centrifuged at 500xg for 5 min at 4°C and resuspended in FACS buffer.

For isolation of immune cells from blood, tubes were prepared with 1 ml of 4% sodium citrate. Blood was transferred to these tubes to prevent clotting and mixed well. 1 ml of FACS buffer was then added. Tubes were underlaid with 1 ml Histopaque (Sigma, 10771-6×100ML) using a Pasteur pipette. Samples were centrifuged at 400xg for 20 min at room temperature. The interface was collected and transferred to a new 15 ml conical tube and washed with 10 ml of FACS buffer to remove Histopaque. After RBC lysis, samples were resuspended in FACS buffer.

For isolation of immune cells from lungs, lungs were removed and collected in a 15-ml centrifuge tube containing 10 ml FACS buffer and placed on ice. Lungs were transferred to a petri dish and macerated with razor blades. 3 ml of lung digestion buffer containing HBSS (Lonza, BE10-508F), 5% FBS, 1 mg/ml collagenase A (Sigma, 2674-500mg), and 0.05 mg/ml DNase I (Roche, 11284932001) was added and transferred to a new 15-ml centrifuge tube. Lungs were incubated at 37°C shaking for 1 hour. In the last 15 minutes of the digestion, lungs were syringed with a 1 ml syringe and 18g needle approximately 20 times. To stop digestion, 10 ml of FACS buffer was added and spun at 400xg for 4 min at 4°C. After lysing RBCs, cells were resuspended in FACS buffer and pushed through a 70-μm cell strainer.

Cavity cells, blood cells and lung single-cell suspensions were washed with PBS and resuspended in PBS containing live/dead fixable aqua dye (Invitrogen, L34957), and incubated at room temperature for 15 min. Cells were then washed and resuspended with FACS buffer. Fc receptors were blocked with anti-CD16/32 Fc block antibody (BD Biosciences, 553142) and Rat IgG then stained with primary antibodies. The antibodies used for staining were as follows: MerTK AF488 (BioLegend, 151504), CD115 PE-CF594 (Biolegend, 135528), I-A/I-E PerCP (BioLegend, 107624), Ly6c PE/Cy7 (BioLegend, 128018), CD64 APC (BioLegend, 139306), CD11b APC/Fire™ 750 (BioLegend, 101262), SiglecF BV421 (BD Biosciences, 565934), CD11c BV605 (117333), F4/80 BV650 (BioLegend, 123149), CD24 PE-CF594 (BD Biosciences, 562477), Ly6G PE (BioLegend, 127607), CD45 Alexa Fluor 700 (BioLegend, 103127). Cells were stained for 30 min at 4°C then washed with FACS buffer. Afterwards, the cells were fixed in 2% paraformaldehyde (PFA) for 20 min at 4°C and resuspended in FACS buffer. Samples were processed on an LSRII (BD Biosciences) or NovoCyte Quanteon (Agilent) and analyzed with FlowJo software (BD Biosciences). For cell sorting, samples were treated similarly, but without fixation, and sorted on a MoFlo Astrios EQ Cell Sorter (Beckman Coulter). FACS buffer used for cell sorting was HBSS without calcium or magnesium and supplemented with 3% FBS, 2mM EDTA and 25mM HEPES. Cells were dispensed into sorting buffer with 4x the amount of FBS.

### Tissue preparation for immunofluorescence and hematoxylin and eosin staining

Mice were sacrificed at the indicated time points. Lungs were inflated with 1 ml of a solution containing 1 part 4% PFA/30% sucrose and 2 parts of optimal cutting temperature compound (OCT) media (Fisher Scientific). Inflated lungs were tied off and stored in 4% PFA/30% sucrose at 4°C overnight. The following day lungs were transferred to 30% sucrose and stored overnight at 4°C. Lungs were then blocked in OCT on dry ice and stored at -80°C. Tissue was sectioned at 10-20 μm on a cryostat. For histological assessment tissues were stained with hematoxylin and eosin. Slides were assessed in a double-blind manner.

For immunofluorescence, slides were mounted with ProLong Glass Antifade Mountant with NucBlue (Invitrogen, P36981). Images were acquired using a Keyence BZ-X710 fluorescent microscope. Image analysis was conducted with two images per section, two sections per slide, two slides per animal with n = 5 animals per condition. Images were analyzed on ImageJ.

### *Ex vivo* labeling of macrophages with PKH26PCL dye

Sorted cell populations were washed twice with serum-free media then resuspended in 500 μl diluent from the PKH26PCL-1KT. In a separate tube, PKH dye solution was prepared at a 5 μM concentration in 500 μl of PKH dye. Dye solution was then added to resuspended cells and incubated for 5 minutes and then washed three times with media containing serum.

### Statistics

All results were presented as the mean ± SEM. R and GraphPad Prism 9.4 were used for statistical analyses. The analyses were conducted using Student’s t test for comparison between two groups and one-way ANOVA was used to test comparisons between multiple groups (*P < 0.05, **P < 0.01, ***P < 0.001, ****P < 0.0001, and ns for not significant).

### Study approval

All experiments with mice were performed in accordance with protocols approved by the University of California Riverside Institutional Animal Care and Use Committee.

## AUTHOR CONTRIBUTIONS

JM conceived and designed the research study. JM, AF, and AN conducted the transcriptomic analyses. IGV and YS provided guidance on DCQ. JM and JPS designed the *in vitro* and *in vivo* experiments. JPS, JM, and SYK conducted the *in vitro* and *in vivo* experiments. JPS and JM analyzed the *in vitro* and *in vivo* experimental data. JM, JPS, MGN, and SYK contributed to data interpretation. JM and JPS wrote the manuscript. All authors read and approved the final manuscript.

## ACKNOWLEDGEMENTS

We thank Maudry Laurent-Rolle, Deborah Spector, Mafalda De Arrabida, Roksana Shirazi, and Xueyan Xu for critical feedback on this manuscript. We thank Adolfo García-Sastre for providing pH1N1 A/California/04/2009 and H1N1 A/Puerto Rico/8/1934 influenza A viruses as well as MDCK cells. The graphical abstract was created with BioRender.com.

