## Supplemental Figures for "Pleural macrophages translocate to the lung during infection to promote improved influenza outcomes"

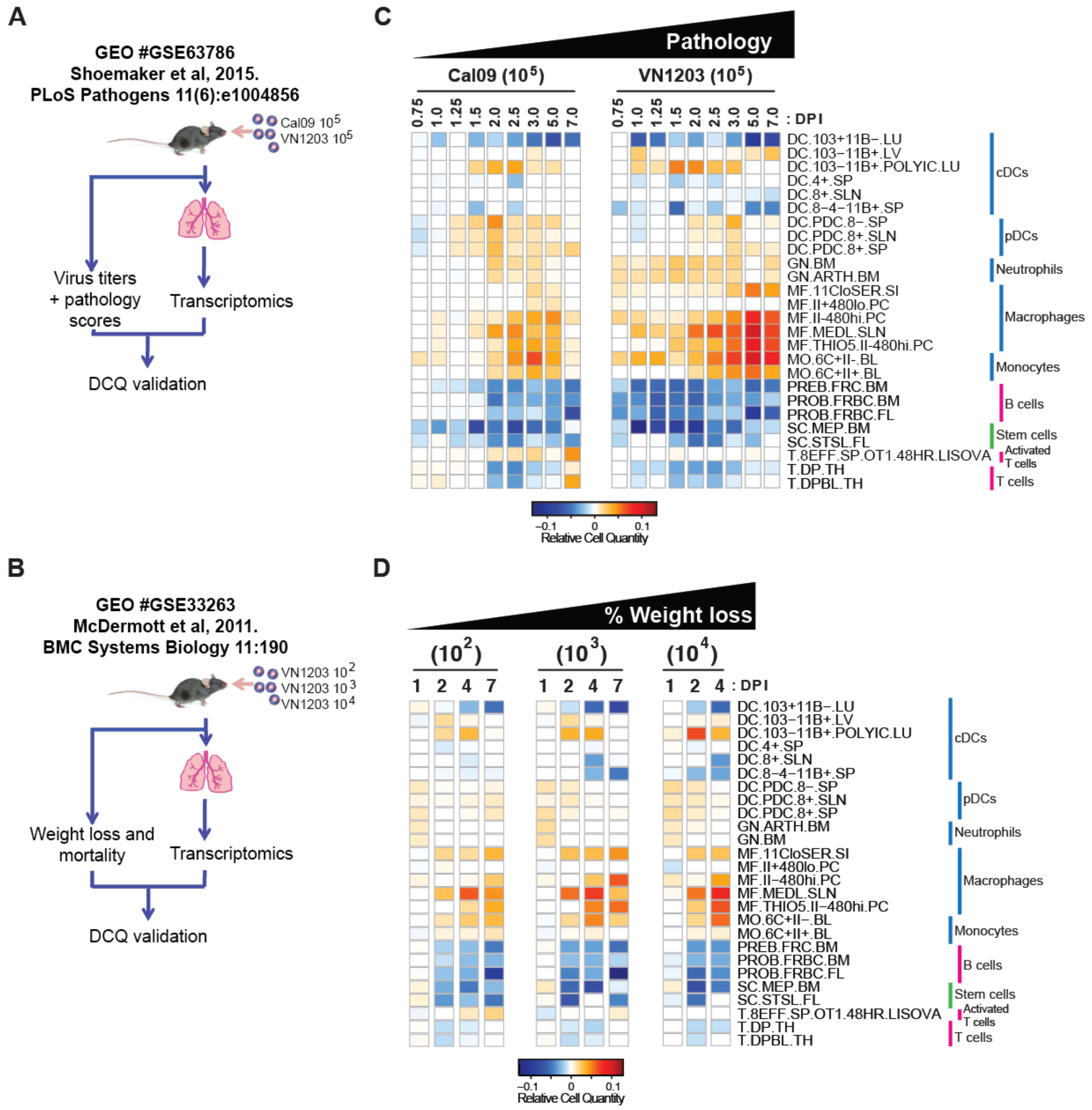


**Figure S1: Immune cell subset predictions hold across additional transcriptomic datasets.** We subjected two C57BL/6 microarray datasets to DCQ (http://dcq.tau.ac.il). (A, B) Schematics showing the experimental design behind the GSE63786 and GSE33263 transcriptomic data. (C, D) Heatmaps for the 26 cell types that were identified as significant in the BALB/c datasets (Figure 2) are shown for the GSE63786 and GSE33263 datasets.

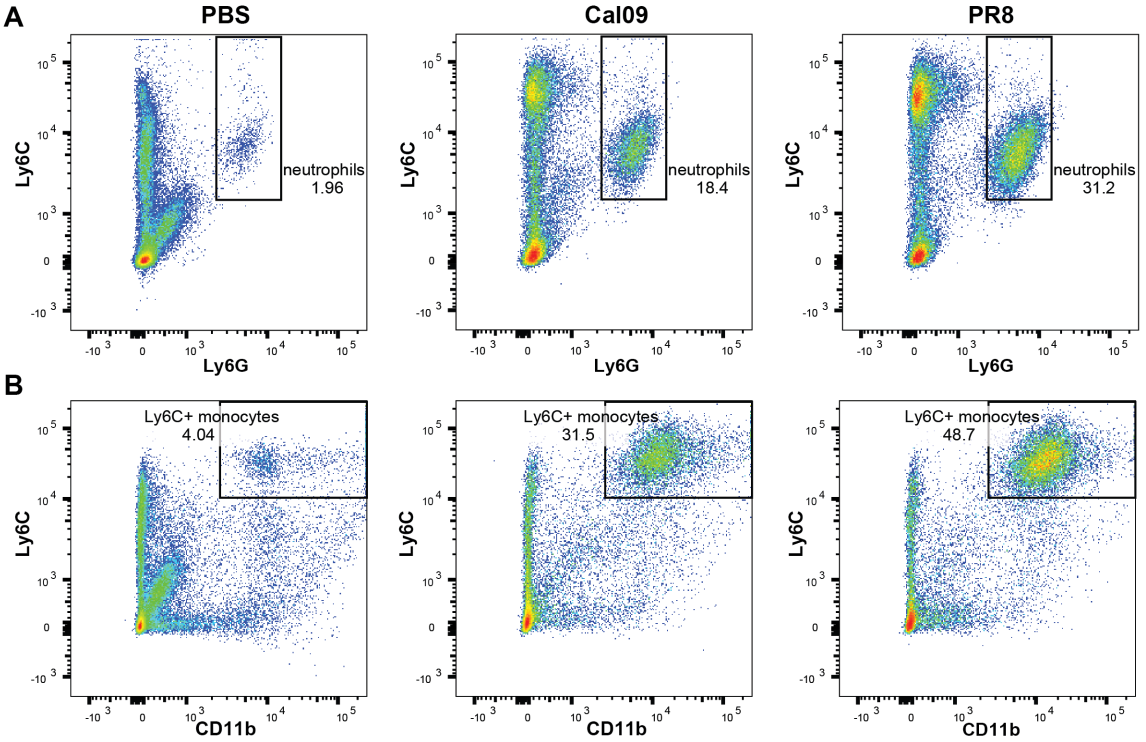


**Figure S2: Flow cytometry validates DCQ predictions.** BALB/c mice were infected intranasally with $10^4$ pH1N1 A/California/09 (Cal09) virus to induce mild disease (n=4) or $10^4$ H1N1 A/Puerto Rico/1934 (PR8) virus (n=4) to induce severe disease. Control mice were mock infected using PBS. On day 3 post infection, lung cells were isolated, stained, and subjected to flow cytometry. Representative plots are shown. (A) Neutrophils = $Ly6C^+Ly6G^+$. (B) $Ly6C^+$ monocytes = $Ly6C^+CD11b^+Ly6G^-$.

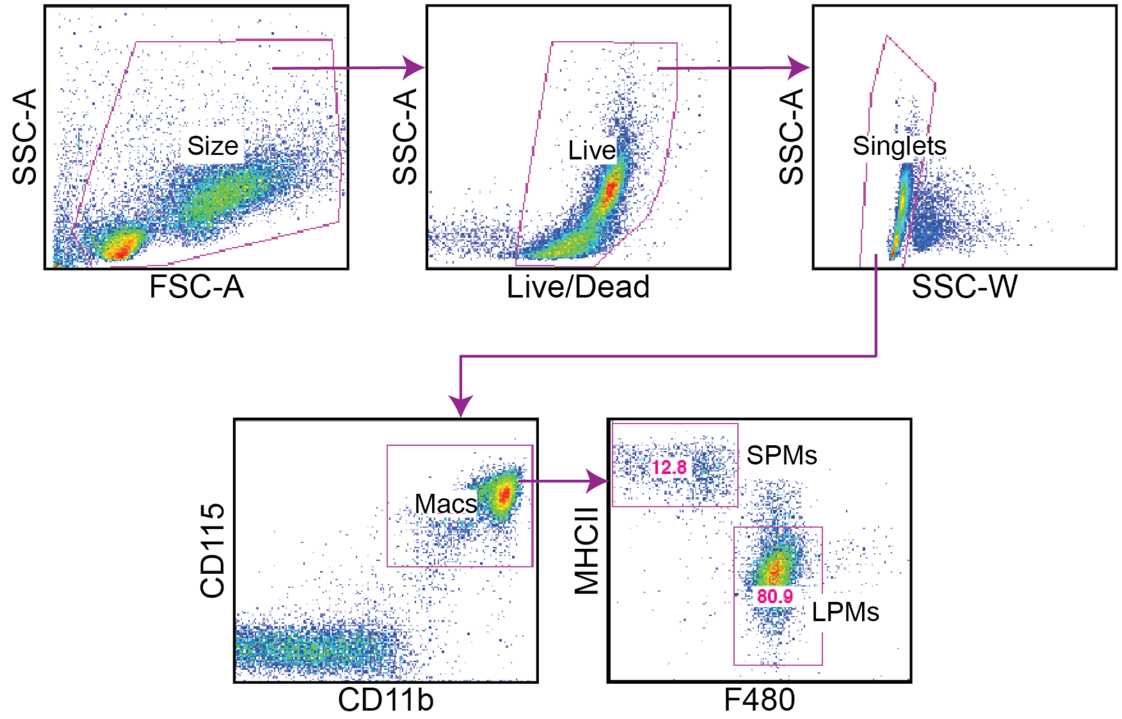


**Figure S3: Flow cytometry gating strategy to identify pleural macrophage subpopulations.** Identification of LPMs and SPMs in the BALB/c pleural fluid following size exclusion, live cells, singlets, and $CD115^{+}CD11b^{+}$ macrophages followed by additional analysis of $F480^{hi}/MHCII^{lo}$ LPMs and $F480^{lo}/MHCII^{hi}$ SPMs.

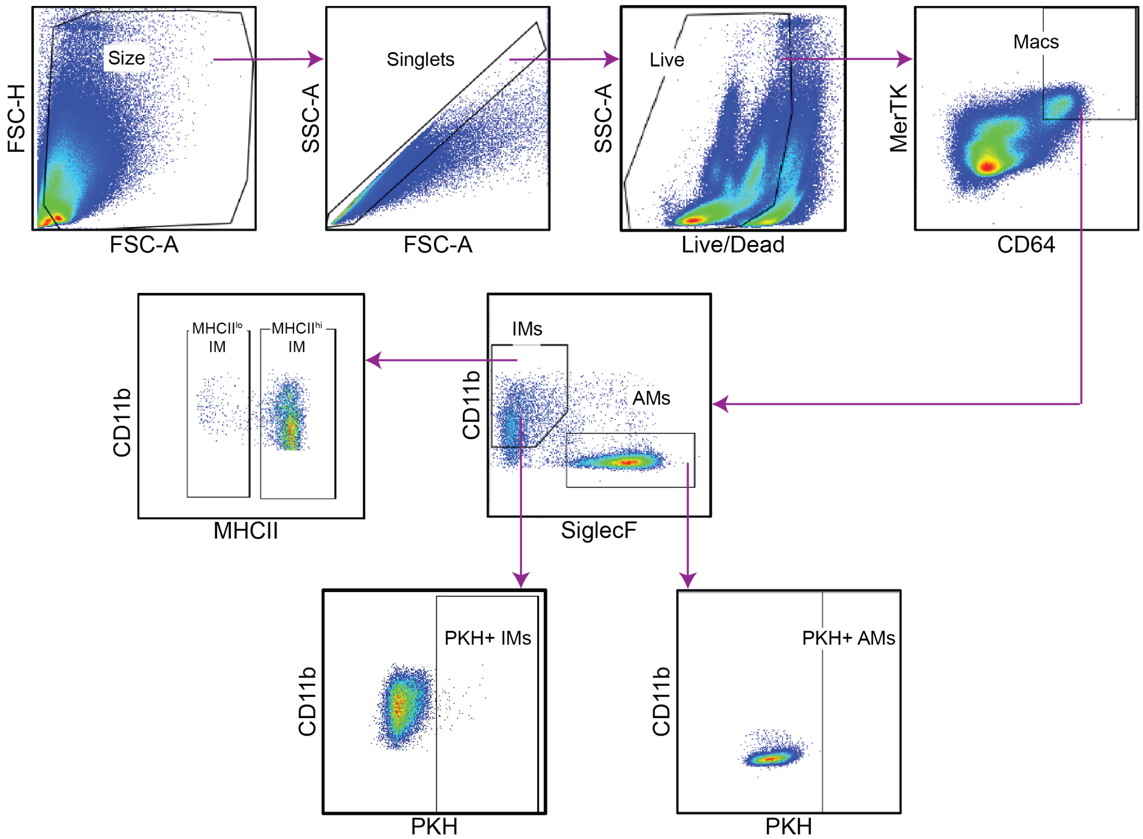


**Figure S4: Flow cytometry gating strategy to identify lung macrophage subpopulations.** Identification of MCHII$^{+}$ and MHCII$^{-}$ IMs, AMs, and PKH$^{+}$ macrophages in BALB/c lung tissue following size exclusion, singlets, live cells, and CD64$^{+}$MerTK$^{+}$ cells followed by additional analysis of PKH on CD11b$^{+}$SiglecF$^{-}$ and MHCII$^{+/-}$ IMs or CD11b$^{-}$SiglecF$^{+}$ AMs.

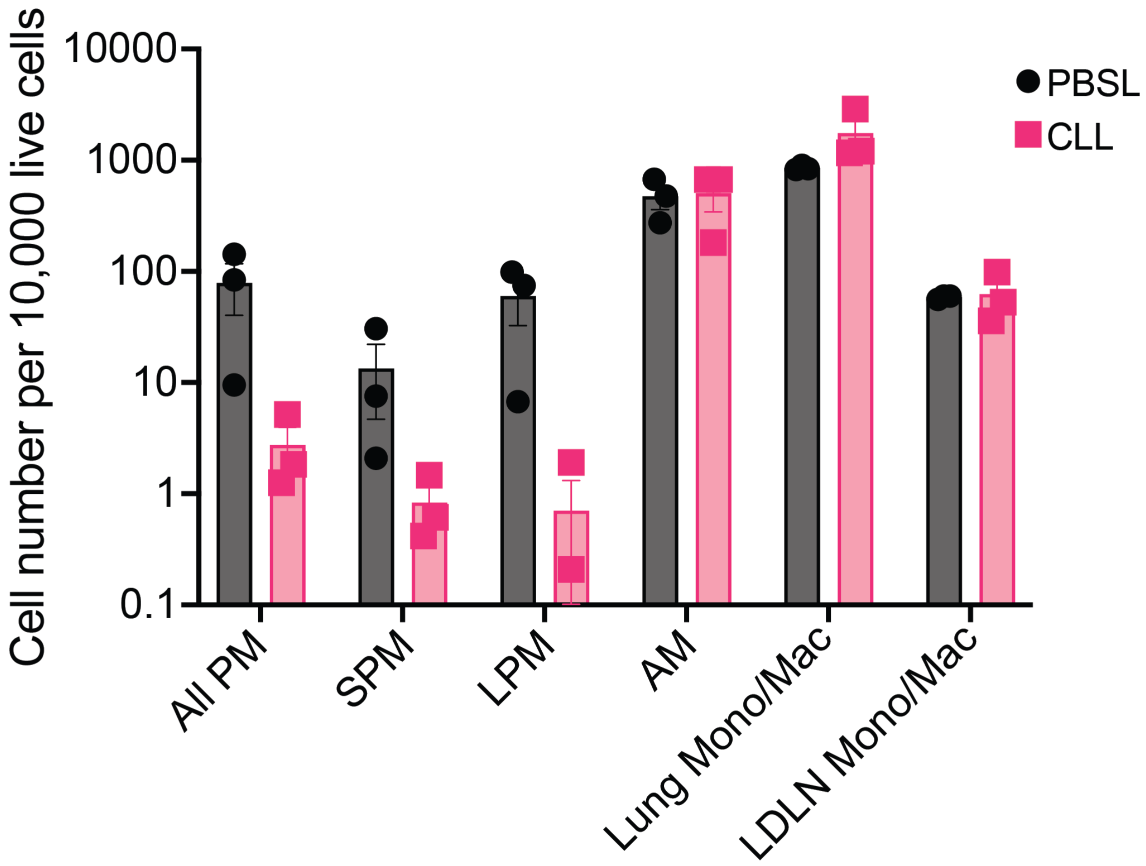


**Figure S5: Intrapleural CLL injection does not impact lung-draining lymph node and lung macrophage populations.** BALB/c mice were injected intrapleurally with CLL or PBS. Pleural fluid, lungs, and lung-draining lymph nodes (LDLN) were isolated 24 hours later for flow cytometry. PM = $CD115^{+}CD11b^{+}$, SPM = $CD115^{+}CD11b^{+}MHCII^{+}F480^{-}$, LPM = $CD115^{+}CD11b^{+}MHCII^{-}F480^{+}$, AM = $CD64^{+}CD11b^{-}SiglecF^{+}$, Other Monocyte/Macrophage (Mono/Mac) = $CD64^{+}CD11b^{+}SiglecF^{-}$.

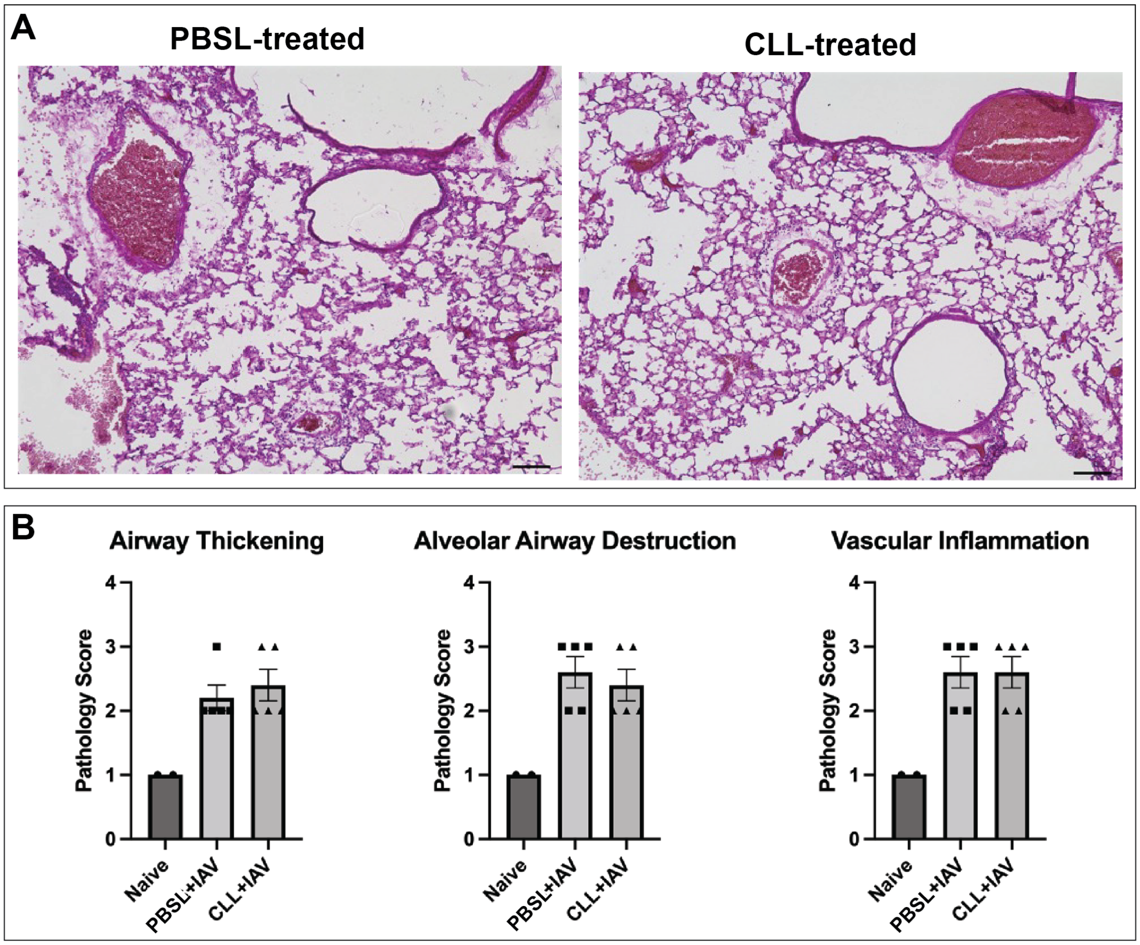


**Figure S6: PM depletion does not affect lung histopathology.** BALB/c mice received an intrapleural injection of CLL or PBSL one day before infection with $10^2$ PFU of Cal09. (A) Representative H&E staining of lungs 9 days post infection in PBSL- or CLL-treated mice (n=4 per group). (B) Quantification of histopathological scores (n=4 per group). Data shown as mean ± SEM (*P<0.05, **P<0.01, ***P<0.001, ****P<0.0001, Student's t-test).

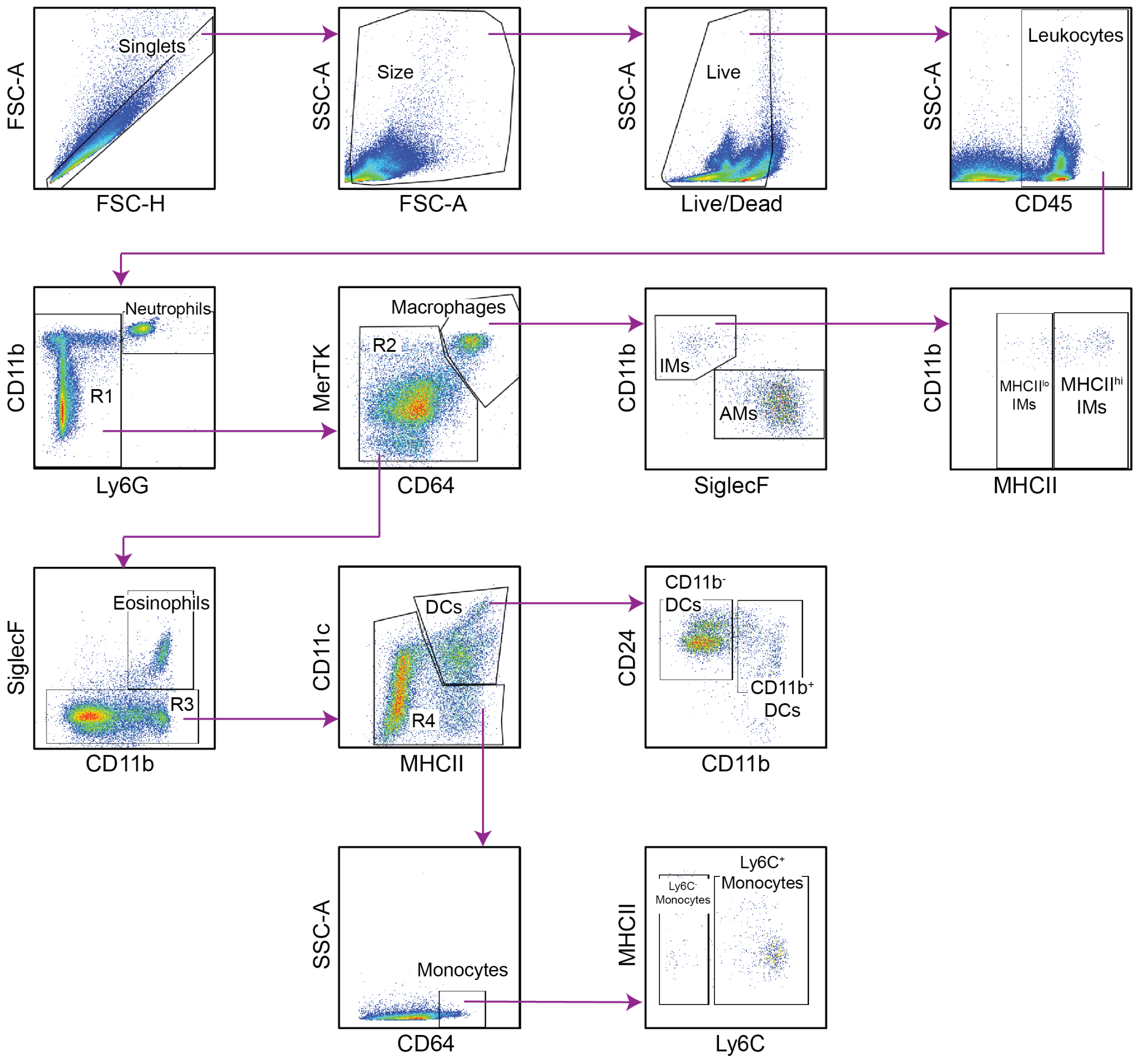


**Figure S7: Flow cytometry gating strategy to identify immune cell subsets.** Identification of immune cell subsets in BALB/c lung tissue following pre-gating on singlets, size exclusion, live cells, and leukocytes ($CD45^+$ cells). Neutrophils were identified as $Ly6G^+CD11b^{hi}$. Macrophages were identified as $Ly6G^-CD64^+Mertk^+$ and divided into IMs ($CD11b^{hi}SiglecF^-$) and AMs ($CD11b^{lo}SiglecF^+$). Two IM subsets were divided based on the expression of $MHCII^{lo/hi}$. Eosinophils were $Ly6G^-CD64^-Mertk^-SiglecF^+CD11b^+$. DCs were $Ly6G^-CD64^-Mertk^-SiglecF^-CD11c^{hi}MHCII^{hi}CD24^{hi}$ and were divided into $CD11b^-$ and $CD11b^+$ DCs. Monocytes were $Ly6G^-Mertk^-$ $SiglecF^-CD24^-$ $CD64^+$ and divided into $Ly6C^-$ or $Ly6C^+$ monocytes. IMs = interstitial macrophages, AMs = alveolar macrophages, DCs = dendritic cells.
